# In-situ Observation of Fast Chloroplast Dynamics in Intact Leaves

**DOI:** 10.1101/2025.10.17.683007

**Authors:** Weihang Geng, Mubin He, Jun Qian

## Abstract

Chloroplast relocation is a hallmark of plant photoprotection, yet has long been regarded as a slow, minute-scale process. Here we overturn this view by directly visualizing chloroplast motility in intact leaves with the second near-infrared (NIR-II, 900–1880 nm) fluorescence confocal microscopy, an optical strategy uniquely suited for deep, in situ imaging. By harnessing the intrinsic long-wavelength fluorescence tail of chloroplasts, this method minimizes scattering, extends imaging depth, and enables simultaneous stimulation-imaging with subcellular resolution—capabilities not attainable with conventional visible fluorescence or multiphoton fluorescence microscopy. Using this approach, we discover that submerged leaves of amphibious plants exhibit remarkably rapid avoidance responses to red light, whereas aerial leaves show negligible relocation. Control experiments exclude influence of the physical structure of leaves, confirming that these contrasting responses reflect distinct physiological adaptations to environmental light regimes. Beyond revealing unexpected speed and flexibility in chloroplast dynamics, our findings establish the second near-infrared fluorescence confocal microscopy as an important tool for direct observation of fast subcellular processes in deep photosynthetic tissues and advancing plant photobiology.

## Introduction

Chloroplasts are dynamic organelles that conduct photosynthesis and can reposition within plant cells to optimize light harvesting under low-light conditions and minimize photodamage under high-light exposure^1,2^. These photorelocation behaviors are primarily regulated by blue and red light, which modulate the polymerization and depolymerization of actin filaments that generate traction forces to drive chloroplast movement^3^. Previous studies using techniques such as transmittance measurement and microscopy have reported that chloroplast relocation typically occurs at a velocity of approximately 1□μm per minute^4–7^. However, existing approaches for observing chloroplast behavior within leaves remain limited. Transmittance measurement methods do not allow for direct, real-time visualization of chloroplast movement^8^. Conventional visible-light microscopy often requires tissue sectioning, which disrupts the in-situ condition^9^. These approaches compromise in situ integrity, temporal resolution, and accuracy of light stimulation during chloroplast behavior observation, which are not conducive to faithfully capture the authentic dynamics of chloroplast movement under natural light conditions, potentially overlooking subtle or transient behavioral features. In natural settings, light intensity may increase abruptly^10^. For plants with complex physiological systems, such as pteridophytes, the relatively slow movement of chloroplasts, if limited to the rates previously reported (1□μm per minute), may be insufficient to prevent light-induced thermal damage or the accumulation of reactive oxygen species (ROS)^11^. Therefore, in situ observation of chloroplast photorelocation dynamics in intact leaves is essential for fully understanding the functional behavior of chloroplasts under physiologically relevant conditions.

To address these challenges, an imaging technique is required that enables precise light stimulation while allowing three-dimensional, in situ, and dynamic observation of chloroplasts in leaves. However, visualizing chloroplasts clearly within intact leaves remains a significant technical challenge. The primary barrier lies in the strong photon scattering of leaf tissues^12,13^, which renders optical observation of deep structures infeasible and severely limits the performance of conventional imaging systems such as wide-field and confocal fluorescence microscopy that use visible light excitation/emission^[13,14]^. Although sample preparation methods such as improved sectioning techniques^15^, vacuum infiltration^16^, and the use of clearing agents enhanced imaging quality^17^, they inevitably disrupted native tissue structures and precluded in situ observation. While two-photon fluorescence microscopy can support in situ imaging of chloroplasts within leaves, its excitation wavelength is not matched with chloroplast absorption peaks, making it unsuitable for simultaneous light stimulation^18^. Applying additional visible light stimulation between imaging intervals impairs real-time tracking, and requires spatial-temporal synchronization between stimulation and imaging^19^. Moreover, femtosecond laser pulses used in two-photon excitation carry high photon energy, which can induce side effects such as phototoxicity and cavitation effects in biological tissues, potentially disturbing the native microenvironment in leaves^20^.

Fluorescence imaging in the second near-infrared window (NIR-II, 900–1880 nm) has emerged as a promising technique for real-time monitoring of living biological specimens^21–23^. In this spectral region, biological tissues exhibit moderate light absorption and reduced photon scattering, which preferentially suppresses scattered photons traveling longer optical paths while allowing ballistic photons with shorter paths to be more effectively detected. This results in improved tissue penetration and superior image clarity for NIR-II fluorescence imaging^24,25^. Taking advantage of these benefits, NIR-II fluorescence imaging has been successfully integrated with a range of existing technologies, including in vivo macroscopic imaging^26,27^, wide-field microscopy^28,29^, light-sheet microscopy^30,31^, and confocal microscopy^32,33^. These combinations have enabled broad applications in image-guided surgery, neuroscience research, tissue visualization, and more. Among them, NIR-II fluorescence confocal microscopy stands out by significantly enhancing imaging depth while preserving intrinsic optical sectioning capability^33,34^, thereby offering great potential for in situ observation of biological dynamics.

In this work, we constructed a NIR-II fluorescence confocal microscope with 671 nm excitation and achieved subcellular-resolution dynamic visualization of rapid chloroplast movements in intact, living plant tissues. Firstly, we found that chloroplasts exhibited strong NIR-II fluorescence and its emission tails could extend to longer than 1200 nm unexpectedly, sufficient for high-quality in situ NIR-II fluorescence microscopic imaging of chloroplasts in living leaves. Then, the focused 671 nm (red) beam functioned as both the stimulation source of chloroplasts in leaves and the excitation source for NIR-II fluorescence imaging simultaneously. Upon continuous scanning, the red-light stimulation triggered a rapid downward migration of chloroplasts in the submerged leaves of amphibious plant along the axial direction within seconds, effectively escaping the illuminated focal plane. This avoidance response was initiated as early as 2.5 seconds after stimulation, with peak velocities reaching up to 1.27□μm/s, approximately 70 times faster than previously reported chloroplast photorelocation movement rates. These findings revealed that chloroplasts could execute photorelocation at unexpectedly high speeds, suggesting that actin filaments can generate stronger traction forces over short time scales. Furthermore, using both *Hydrocotyle sibthorpioides* and *Hygrophila pinnatifida*, we found that chloroplasts in submerged and aerial leaves of the same amphibious plant responded differently to red-light stimulation. Chloroplasts in submerged leaves rapidly avoided the illumination zone, whereas those in aerial leaves did not exhibit such avoidance behavior. It is attributed to that submerged and aerial leaves are exposed to different red-light environments, since red-light is easily absorbed by water and cannot penetrate deep inside water. These results underscore the crucial role of environmental context in shaping chloroplast photoreactivity, which is supported by the optical principle, that amphibious plants adopt distinct photosynthetic strategies adapted to the respective habitats of leaves.

## Results

### Optimization of the excitation/emission wavelength for fluorescence confocal microscopic imaging of chloroplasts within intact leaves

To optimize the imaging quality of intact leaves, we first investigated the excitation wavelengths of the fluorescence confocal microscope. The absorption and fluorescence spectra of chloroplasts are shown in Figure 1a. Two prominent absorption peaks were observed at 438 nm and 678 nm, corresponding to excitation wavelengths that can effectively stimulate the intrinsic fluorescence of chloroplasts. Optical sectioning images of intact leaf samples were then acquired using a fluorescence confocal microscopic system excited by 405 nm and 640 nm lasers. In theory, the shorter-wavelength 405 nm excitation should provide substantially higher spatial resolution than the longer-wavelength 640 nm excitation^35^. However, as shown in Figures 1b and 1c, strong photon scattering within the leaf tissues led to the opposite result at a depth of 15 μm. Under 640 nm excitation, the internal structure of chloroplasts with 564 nm width could be clearly resolved, whereas the same structure could not be detected under 405 nm excitation. Due to reduced scattering and attenuation, the longer-wavelength 640 nm excitation beam (Figure 1d) was able to focus more tightly and penetrate deeper into the leaf tissues, resulting in confocal images with significantly improved spatial resolution and SBR. Nevertheless, fluorescence confocal microscopy excited at 640 nm still exhibited limitations. As shown in Figure 1e, images obtained at shallow depths displayed pronounced structural overlap (highlighted by the white dashed boxes). At an imaging depth of 30 μm, the SBR dropped sharply, and image contrast was nearly lost.

**FIGURE 1.**
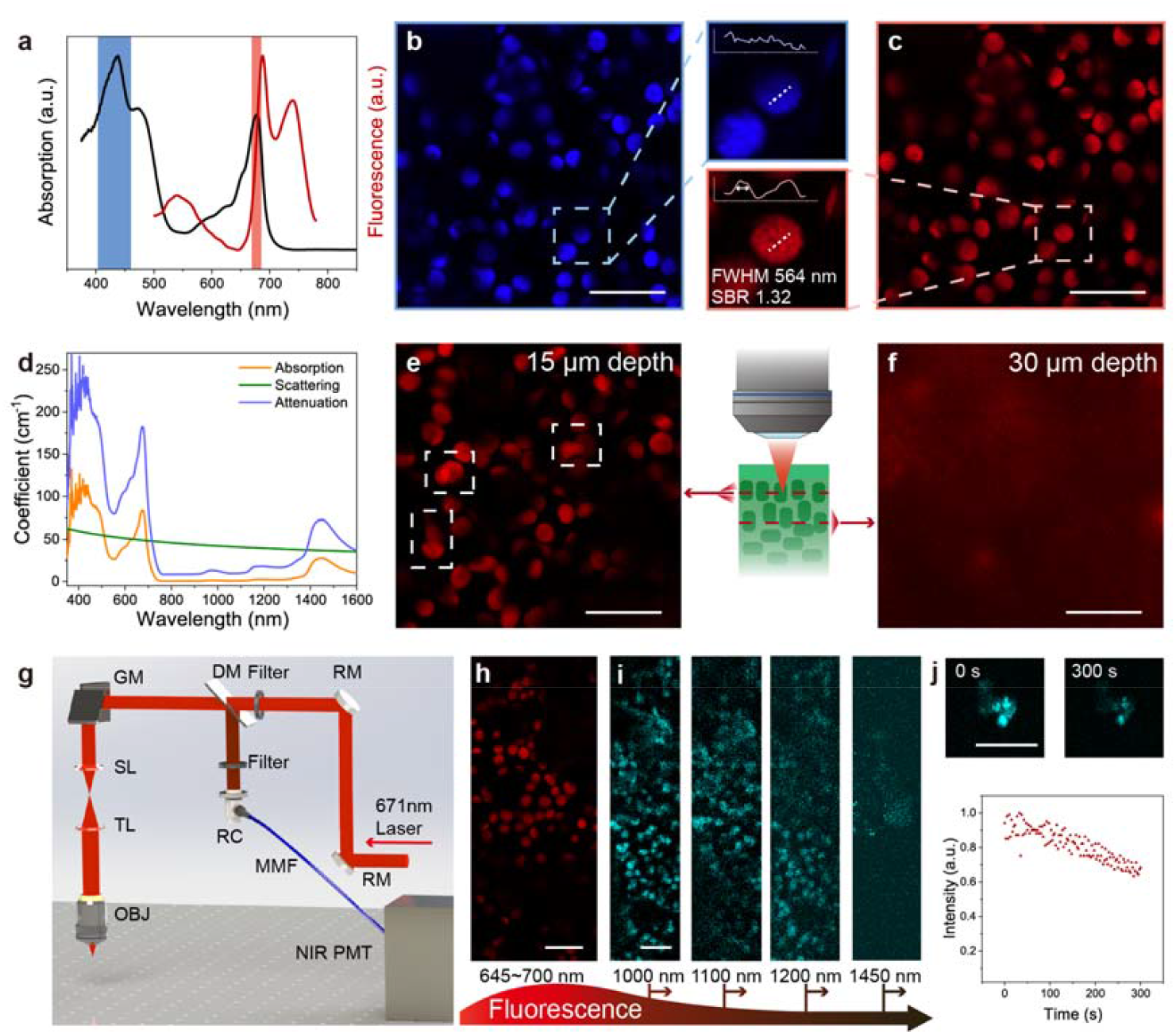
Fluorescence confocal microscopic imaging of chloroplasts within intact leaves, based on various excitation and emission wavelengths. (a) Absorption and fluorescence spectra of chloroplasts, showing two major absorption peaks (∼438 nm and ∼678 nm) in the blue and red spectral regions. (b, c) Fluorescence confocal microscopic images of chloroplasts inside intact leaf under excitation of 405 nm (c) and 640 nm (d) lasers. Spectral range of detected fluorescence: 645-700 nm. Inset shows magnified areas and the intensity curves marked by the dotted lines. The full width at half maximum (FWHM) and signal-to-background ratio (SBR) of the signal peak indicated by the double sided arrow is presented. (d) Dependence of leaves’ absorption coefficient, scattering coefficient and effective attenuation coefficient on wavelength. (e) Fluorescence confocal microscopic image at 15 μm depth under excitation of 640 nm laser, revealing overlapping distributions of chloroplasts across different depths in certain regions (indicated by the white dashed boxes). Spectral range of detected fluorescence: 645-700 nm. (f) Fluorescence confocal microscopic image at 30 μm depth under excitation of 640 nm laser, showing sharp degradation in SBR and loss of image contrast. Spectral range of detected fluorescence: 645-700 nm. (g) Schematic diagram of the NIR-II fluorescence confocal microscope, where fluorescence is collected by a NIR detector after passing through a long-pass filtering with various cutoff wavelength. Abbreviations: RM (Reflecting Mirror), DM (Dichroic Mirror), GM (Galvanometers), SL (Scan Lens), TL (Tube Lens), RC (Reflective Collimator), OBJ (Objective Lens), MMF (Multi-mode Fiber), NIR PMT (NIR sensitive Photomultiplier Tube). (h) Fluorescence confocal microscopic image of chloroplasts in a living leaf, with detected spectral range of 645-700 nm. Excitation wavelength: 640 nm. (i) NIR-II fluorescence confocal microscopic images of chloroplasts in a living leaf, with detected spectral range beyond 1000 nm, 1100 nm, 1200 nm and 1450 nm. Excitation wavelength: 671 nm. (j) Time-resolved NIR-II fluorescence (beyond 1000 nm) intensity curve under continuous scanning of 671 nm laser. Power in front of the objective: 1 mW. Insets show images at start (0 s) and end (300 s) time points. Scale bar: 20□μm.

Since the excitation wavelength of chloroplasts is relatively difficult to modify, we aimed to enhance their imaging depth and contrast in intact leaves by optimizing the fluorescence detection band instead. As shown in Figure 1a, chloroplasts exhibit a peak emission at 687 nm. In our previous studies, we demonstrated that methylene blue and near-infrared fluorescent proteins, with peak emissions above 700 nm, possess long fluorescence tails extending beyond 900 nm, which are advantageous for deep-tissue and high-contrast NIR-II imaging^36,37^. To determine whether chloroplasts also exhibit sufficiently strong tail emission in the NIR-II region, we constructed an NIR-II fluorescence confocal microscopic system (Figure 1g), using a 671 nm excitation laser-close to the chloroplast absorption peak at 678 nm-for validation. As shown in Figure 1i, even at a power as low as 1 mW in front of the objective, bright fluorescence signals were detectable beyond 1000 nm. Remarkably, emission beyond 1200 nm remained observable under confocal microscopy, and the fluorescence tail was still distinguishable beyond 1450 nm. Furthermore, Figure 1j shows that the NIR-II fluorescence intensity of chloroplasts decreased by only 40% after 300 s of continuous 671 nm laser illumination, indicating a certain degree of photobleaching resistance.

### Deep-tissue three-dimensional imaging of chloroplasts in living leaves using NIR-II fluorescence confocal microscopy

We next employed the 671 nm-excited NIR-II fluorescence confocal microscope to image living leaves of *Hydrocotyle sibthorpioides*. As shown in Figure 2a-c, chloroplasts were clearly distinguishable at a depth of 15 μm. At 30 μm, their structural details remained sharp, in stark contrast to the results obtained using a 640 nm-excited visible fluorescence confocal microscope (Figure 1f), where imaging contrast was completely lost due to strong scattering of visible fluorescence within leaf tissues. Remarkably, even at a depth of 45 μm, chloroplasts could still be optically resolved with the NIR-II fluorescence confocal microscope. Three-dimensional reconstruction (Figure 2d, e) further revealed the spatial distribution of chloroplasts at depths between 20 and 40 μm within the leaf. Owing to the inherently lower axial resolution of confocal microscopy compared to its lateral resolution, the NIR-II fluorescence signals exhibited more pronounced axial tailing. Nevertheless, the average size of chloroplasts could still be reliably estimated by measuring their median position of the fluorescent tailing (Figure 2f). Taken together, these results demonstrate that the strong fluorescence of chloroplasts beyond 1000 nm enables their clear visualization deep within living leaves using NIR-II fluorescence confocal microscopy.

**FIGURE 2.**
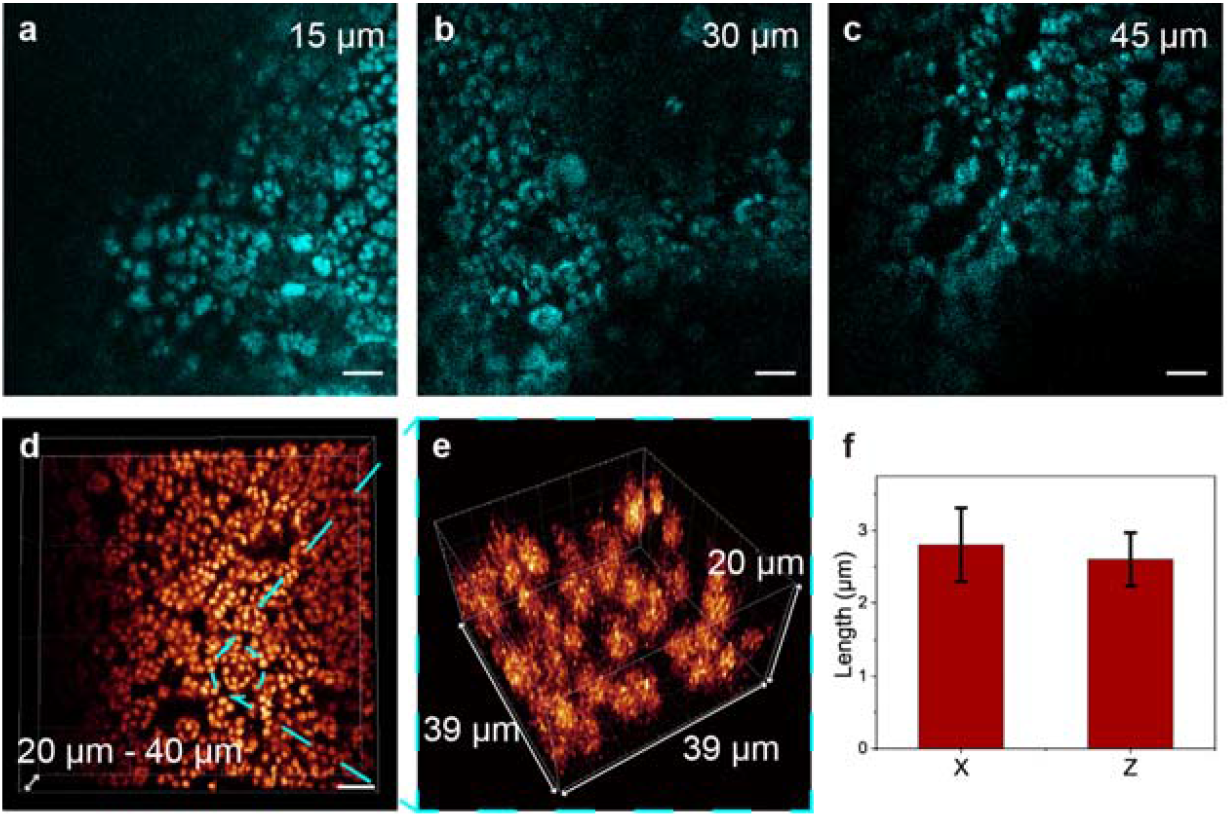
NIR-II fluorescence confocal microscopic imaging of chloroplasts within intact leaves of Hydrocotyle sibthorpioides. (a-c) NIR-II fluorescence confocal microscopic images of chloroplasts at depth of 15 μm, 30 μm and 45 μm within intact leaves, under excitation of 671 nm laser with detection beyond 1000 nm. Power in front of the objective: 1 mW. (d) Three-dimensional reconstruction of NIR-II fluorescence confocal microscopic images of chloroplasts at depths of 20-40□μm within the leaf. (e) Magnified view of the region selected in (d), showing the spatial distribution of chloroplasts in a more detailed manner. (f) Measurement of the diameters in the axial and lateral directions of 10 chloroplasts in (e), obtaining the average size of chloroplast in *Hydrocotyle sibthorpioides*. Scale bars: 20 μm.

### In Situ Dynamic Imaging of Rapid Chloroplast Avoidance Movement under Red Light Stimulation

To achieve dynamic in situ observation of chloroplasts within intact leaves, we used NIR-II fluorescence confocal microscope to image living submerged leaves of *Hydrocotyle sibthorpioides*. The 671 nm continuous-wave (CW) laser in the microscopic system simultaneously served as the excitation and the light stimulus sources for chloroplasts. After 20 s of focused 671 nm laser scanning at a depth of ∼30 μm, the distribution of chloroplast fluorescence within the imaged area changed markedly. Chloroplasts in the region outlined in Figure 3a exhibited lateral migration and axial displacement, as shown in Figure 3b. The observed fluorescence defocusing indicated chloroplast displacement away from the original focal plane along the z-axis. To validate these movements, we performed three-dimensional reconstructions of chloroplast positions before and after continuous laser stimulation (Figures 3c–g). Figure 3c-e showed three-dimensional reconstruction of the two target chloroplasts within a range of ±15 μm of the scanning depth after their movements. Comparisons of chloroplast positions before and after red light stimulation revealed both lateral migration and axial displacement to larger depths. Figure 3e indicated that the centroid of chloroplast fluorescence shifted ∼5 μm deeper within just 20 s. Chloroplasts within the imaging field of view exhibited collective downward migration (Figure 3g), indicating a clear red-light avoidance response in submerged *Hydrocotyle sibthorpioides* leaves. These findings indicated that chloroplasts in *Hydrocotyle sibthorpioides* can rapidly reposition within seconds to avoid red-light exposure.

**FIGURE 3.**
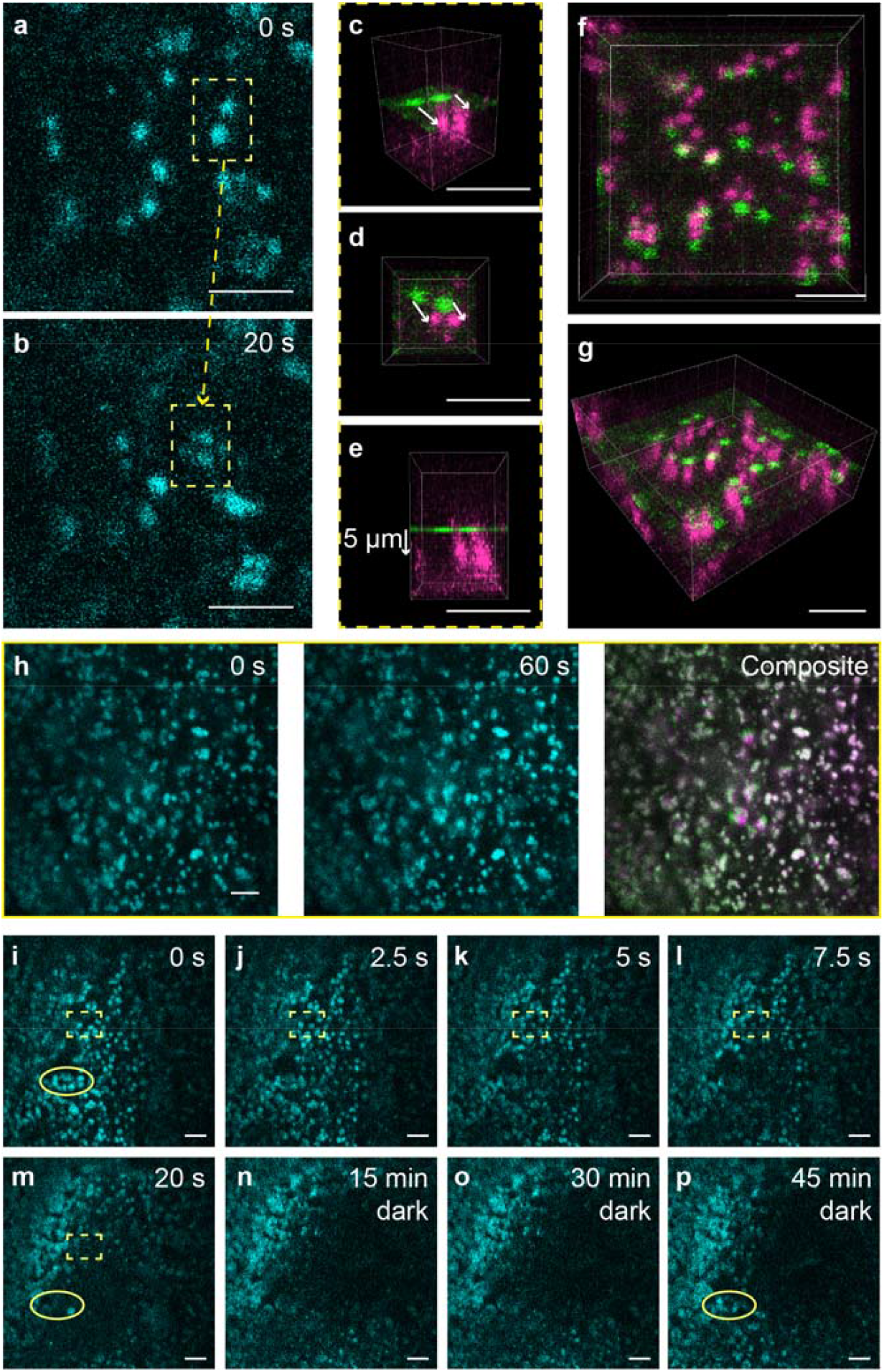
NIR-II fluorescence confocal microscopic imaging of chloroplast within intact leaves of *Hydrocotyle sibthorpioides* under 671 nm laser excitation and stimulation with detection beyond 1000 nm. Power in front of the objective: 1 mW. (a, b) In situ NIR-II fluorescence confocal microscopic images of chloroplasts within an intact living leaf at around 30 μm depth, before (a) and after (b) continuous scanning for 20 seconds under 671 nm laser. Yellow dashed boxes highlight two chloroplasts that, while initially positioned within the focal plane, displayed pronounced three-dimensional relocation during scanning. (c–e) Dual-color three-dimensional reconstruction within the yellow dashed boxes at (a, b) shows single-layer imaging at the scanning depth before scanning (green) and reconstruction at ±15 μm range of the scanning depth after scanning (magenta): (c) side view, (d) top view, and (e) front view, demonstrating spatial tracking of the two displaced chloroplasts. (f, g) Whole-field dual-color three-dimensional reconstruction under the same conditions as (c-e), showing general chloroplasts movements after scanning. (h) Ex-situ NIR-II fluorescence confocal microscopic images of intact metabolically inactive leaf before (green) and after (magenta) continuous scanning for 60 seconds under 671 nm laser. The right image is their merged one. (i-m) Time-lapse in situ NIR-II fluorescence confocal microscopic images of chloroplasts within an intact living leaf under continuous scanning of 671 nm laser. The dashed boxes highlight chloroplast avoidance movements along z-axis. (n-p) NIR-II fluorescence confocal microscopic images of chloroplasts within an intact living leaf under dark adaptation after light stimulation of (i-m). The solid line ellipses show chloroplasts repositioning. Scale bars: 20 μm.

To confirm that this behavior reflected an active physiological response, control experiments were conducted on metabolically inactive leaves (detached and kept at room temperature for two hours). In Figure 3h, chloroplasts in metabolically inactive leaves remained stationary throughout continuous 671 nm laser scanning. Dual-color images acquired before (0 s) and after (60 s) red-light stimulation showed high spatial overlap, effectively excluding thermal expansion or pressure-driven swelling as confounding factors. This strongly supports the conclusion that light-induced chloroplast migration in living leaves originates from active intracellular physiological mechanisms.

Real-time imaging at 2.5 s intervals during continuous 671 nm laser scanning (Figures 3i–m) captured the dynamic process of rapid chloroplast migration in live leaves. Chloroplasts responded as early as the first acquired frame (2.5 s). Within 20 s, chloroplast fluorescence at the field center (indicated by the dashed boxes) gradually diminished and disappeared, indicating displacement away from the focal plane along the z-axis. This dynamic process is illustrated in Video S1. After dark adaptation, chloroplasts gradually returned to their original uniform distribution (Figures 3n–p). Previously vacated regions, as the solid line ellipses shown in Figures 3i, 3m and 3p, were reoccupied by chloroplasts. To our knowledge, this represents the first direct *in situ* demonstration of rapid, dynamic, light-responsive chloroplast behavior within intact leaves, underscoring their remarkable adaptability to fluctuating light conditions.

### Differences of Chloroplast Photorelocation Response in Aerial and Submerged Leaves of Amphibious Plants

Due to differential absorption of various spectral bands of sunlight by water, submerged leaves and aerial leaves experience distinct light environments. The leaves of amphibious plants exhibit adaptive photosynthetic mechanisms in diverse environments^38,39^, which extend to differences in chloroplast photorelocation movement. The amphibious plant *Hydrocotyle sibthorpioides* commonly inhabits moist soil or submerged aquatic environments. As shown in Figure 4a, upon full submersion of water, aerially grown leaf of *Hydrocotyle sibthorpioides* undergoes senescence and abscission, followed by regeneration of submerged-adapted leaves over several days. Using NIR-II fluorescence confocal microscopic imaging, we observed in situ distinct chloroplast responses between aerial and submerged leaves of same plants under red light stimulation with the same power and stimulation time. As shown in Figures 4b and 4c, under continuous scanning of 671 nm laser, submerged leaves exhibited clear chloroplast avoidance movement, in contrast to the negligible response observed in aerial leaves. Instead, increase and diffusion of chloroplast fluorescence signals were observed in aerial leaves. To validate these findings, we conducted analogous experiments on another amphibious species, *Hygrophila pinnatifida* (Figures 4d, e), which produced consistent results. Dual-color images (Figures 4f-i) and the quantitative changes in fluorescence intensity (Figures 4n-q) further highlight these differences, namely, a substantial decrease in the fluorescence intensity of active submerged leaves and a significant increase in that of active aerial leaves following 50 s continuous red-light stimulation.

**FIGURE 4.**
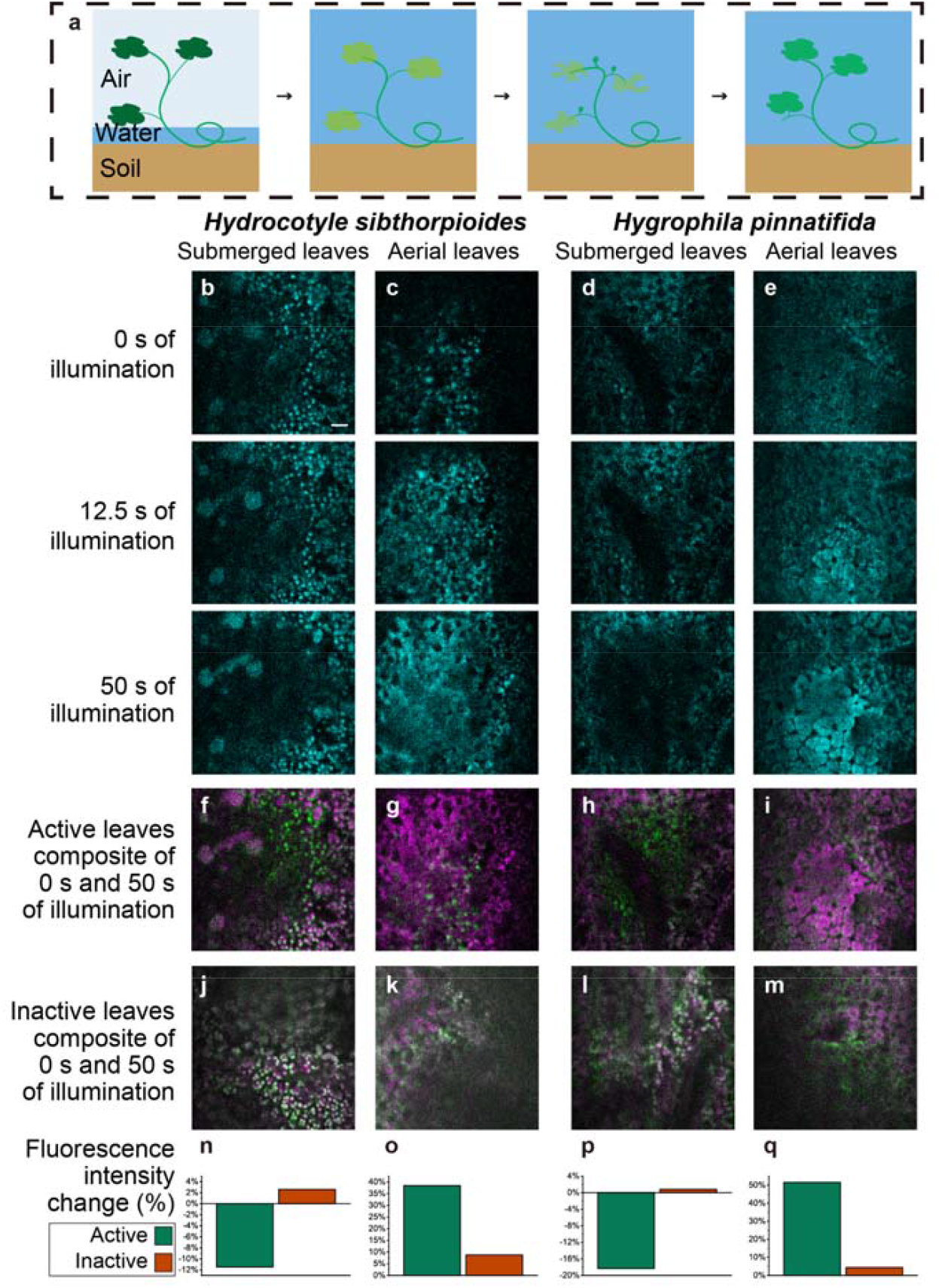
Photorelocation responses of chloroplasts in aerial and submerged leaves of two amphibious plant species to red light. (a) Schematic illustration of the transition from aerial to submerged leaves in two amphibious plants: *Hydrocotyle sibthorpioides* and *Hygrophila pinnatifida*. Plants that initially develop in air undergo leaf yellowing, dissolution, and abscission upon submersion of water, followed by the regeneration of submerged-adapted new leaves. (b–e) In situ time-lapse NIR-II fluorescence confocal microscopic images of chloroplasts in submerged and aerial leaves of two species of amphibious plants, under 50 s continuous scanning of 671 nm laser with detection beyond 1000 nm. Power in front of the objective: 1 mW. (f–i) Dual-color merged images of chloroplast distributions at 0□s (green) and 50□s (magenta) across different leaf types. (f), (g), (h) and (i) correspond to (b), (c), (d) and (e), respectively. (j–m) Dual-color merged NIR-II fluorescence confocal microscopic images of chloroplast distributions in intact metabolically inactive leaves merged at 0□s (green) and 50□s (magenta) under the same illumination conditions of (b-e). (n-q) The ratio of change in chloroplast fluorescence intensity between 50 s and 0 s to the chloroplast fluorescence intensity at 0 s across different leaf types (aerial and submerged) and physiological states (metabolically active and inactive samples). Green and orange values denote the changes in fluorescence intensity of active and inactive leaves, respectively, under the leaf types of the corresponding column. Scale bars: 20 μm.

To verify that these behaviors were physiological, intact metabolically inactive leaves (detached and kept at room temperature for two hours) were imaged by NIR-II fluorescence confocal microscope. In Figures 4j-m, chloroplasts in inactive aerial and submerged leaves exhibited minimal displacement or fluorescence change under the same light stimulation. The movement speed of chloroplasts was slow, similar to the previous conclusion^4^, which indicated the significance of in situ observation. The quantitative changes of fluorescence intensity of metabolically inactive leaves (Figure 4n-q) were not distinct compared to that of the living (metabolically active) leaves. This confirmed that the responses observed in situ in living leaves reflect active physiological behavior rather than physical artifacts, including thermal damage, optical aberrations, or mechanical drift.

These results indicated that, relative to aerial leaves, submerged leaves exhibit heightened sensitivity to red light and an enhanced capacity for rapid avoidance under strong red-light stimulation in *Hydrocotyle sibthorpioides* and *Hygrophila pinnatifida*. It suggested that the adaptation of amphibious plants to distinct light environments is mirrored in their characteristic patterns of chloroplast photorelocation. The observations provided direct visual evidence of these adaptations, enhancing understanding of the complex regulation of plant photoresponses.

Furthermore, we proposed a possible optical basis for this mechanism. Considering the stronger absorption of red light by water, submerged leaves develop under comparatively low red-light conditions. In contrast, the aerial leaves inhabit an environment characterized by abundant red-light exposure, to which they have gradually adapted. This may underlie the observed heightened sensitivity to red light of submerged leaves, which potentially serves to mitigate photodamage under identical red-light stimulation as the aerial leaves^40^. From another perspective, in comparison to red light, water exhibits lower absorption of blue light. Consequently, the submerged leaves are situated in an environment with ample blue light exposure, indicating their adaptation to such illumination conditions. Supporting this view, Video S2 demonstrated that chloroplasts exhibit minimal movement, limited to minor rotation, following 3 minutes of continuous higher-power blue-light excitation. This contrasted sharply with the rapid avoidance movements observed under red-light stimulation.

### Control Experiments to Verify the Distinct Chloroplast Photoresponse in Amphibious Plants

To rule out the possibility that differences in chloroplast photoresponse under the same scanning laser power were caused by variation in light absorption or scattering between leaf types, which could alter the effective irradiance at mesophyll cells, we conducted a series of control experiments on *Hydrocotyle sibthorpioides*. Adjusting the power of red-light illumination during imaging of submerged leaves never reproduced the widespread fluorescence enhancement and diffusion seen in aerial leaves. Under continuous red-light illumination, submerged leaves consistently failed to mimic responses of aerial leaves: even at irradiances exceeding the photodamage threshold, chloroplasts maintained a robust avoidance behavior (Figure 5a), whereas at lower laser power they exhibited localized aggregating toward the laser focused plane (Figure 5b). In contrast, even under high-power red-light illumination sufficient to induce photodamage, chloroplasts in aerial leaves showed no detectable avoidance (Figure 5c). These experiments confirm that the rapid photoresponse of chloroplasts in submerged leaves are genuine physiological traits and exclude differences in tissue scattering or light penetration at comparable depths as the primary cause, underscoring the distinctiveness of the chloroplast avoidance mechanism in submerged leaves of amphibious plants.

**FIGURE 5.**
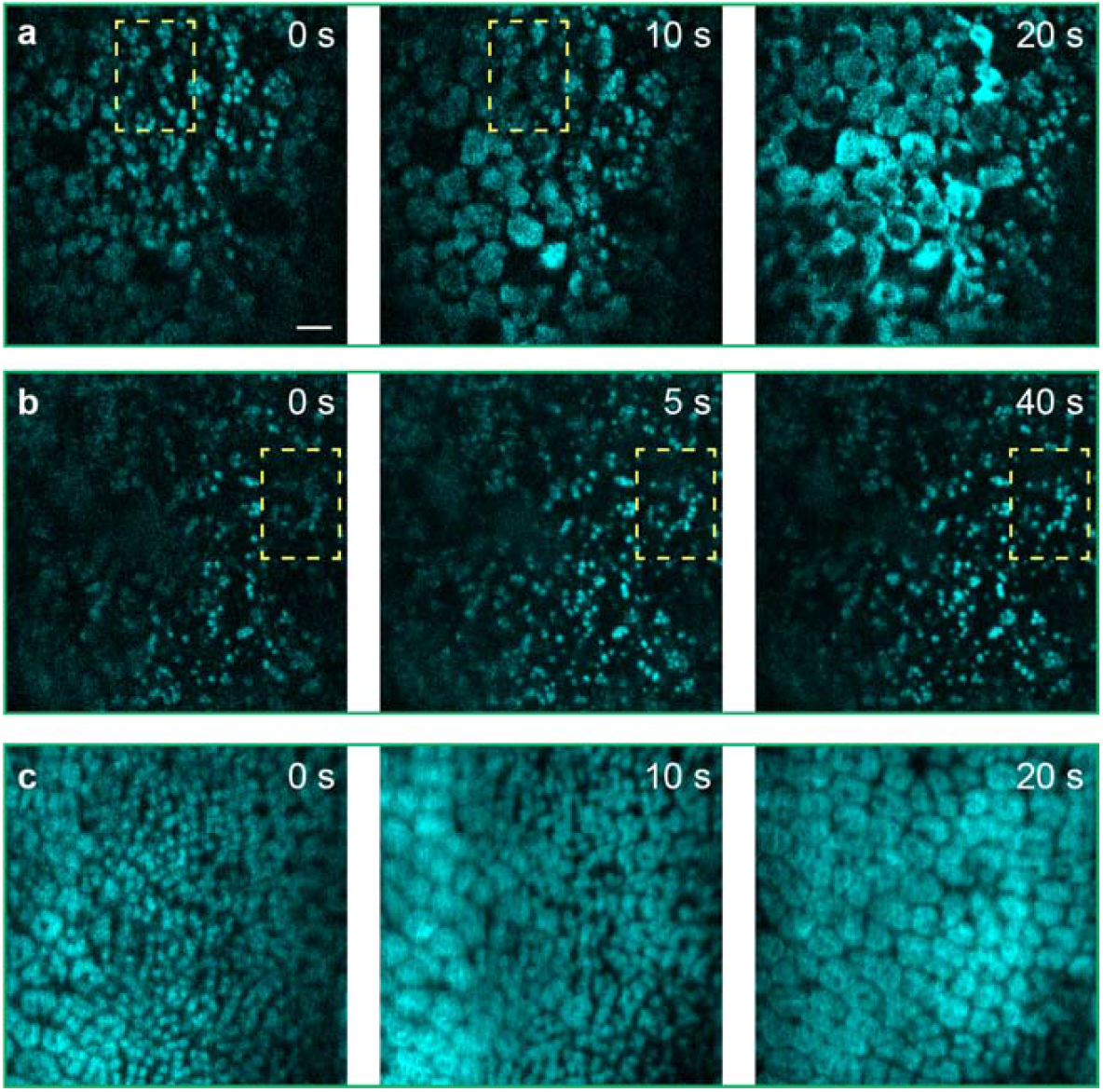
In situ NIR-II fluorescence confocal microscopic images of divergent chloroplast photoresponses in submerged and aerial leaves of *Hydrocotyle sibthorpioides* under 671 nm laser excitation with detection beyond 1000 nm. (a) Submerged leaves still exhibited chloroplast avoidance movement (highlighted by the dashed boxes) under continuous scanning of high-power laser sufficient to induce photodamage (image at 20 s showed obvious photodamage). Power in front of the objective: 4.5 mW. (b) Chloroplasts in submerged leaves aggregated to the focal plane (highlighted by the dashed boxes) under continuous scanning of low-power laser. Power in front of the objective: 0.2 mW. (c) Aerial leaves did not exhibit chloroplast avoidance movement under continuous scanning of high-power laser sufficient to induce photodamage. Power in front of the objective: 4.5 mW. Scale bars: 20 μm.

### Time-lapse Quantification of Rapid Chloroplast Photoresponses Dynamics

Compared to previously reported chloroplast migration velocities of only a few micrometers per minute, our measurements revealed a significantly accelerated chloroplast avoidance response under red-light stimulation. We therefore quantitatively measured the migration trajectories and velocities of chloroplasts in live leaves during light stimulation. Given the temporal constraints of volumetric imaging, we prioritized our analysis of XY plane displacement, providing a conservative estimate of true 3D velocities. As shown in Figure 6a-c, continuous confocal microscopic imaging at 2.5-second intervals enabled tracking of both lateral displacement and axial defocusing behavior of chloroplasts. Some chloroplasts exhibited lateral movement in conjunction with axial (Z-direction) displacement. Analysis of lateral displacement across multiple chloroplasts revealed a maximum in-plane velocity of 0.84 μm/s (Figure 6d-e). Furthermore, samples with reduced metabolic activity (detached and kept at room temperature for ∼30 minutes) could exhibit substantial lateral displacement without significant axial movement (Figure 6f-h). This observation enabled the assessment of chloroplast lateral movement to effectively reflect the intrinsic chloroplast movement velocity. Peak velocity of the rapid chloroplast photoresponse reached up to 1.27 μm/s (Figure 6i-j), demonstrating an order of magnitude higher than previous reports^4–7^.

**FIGURE 6.**
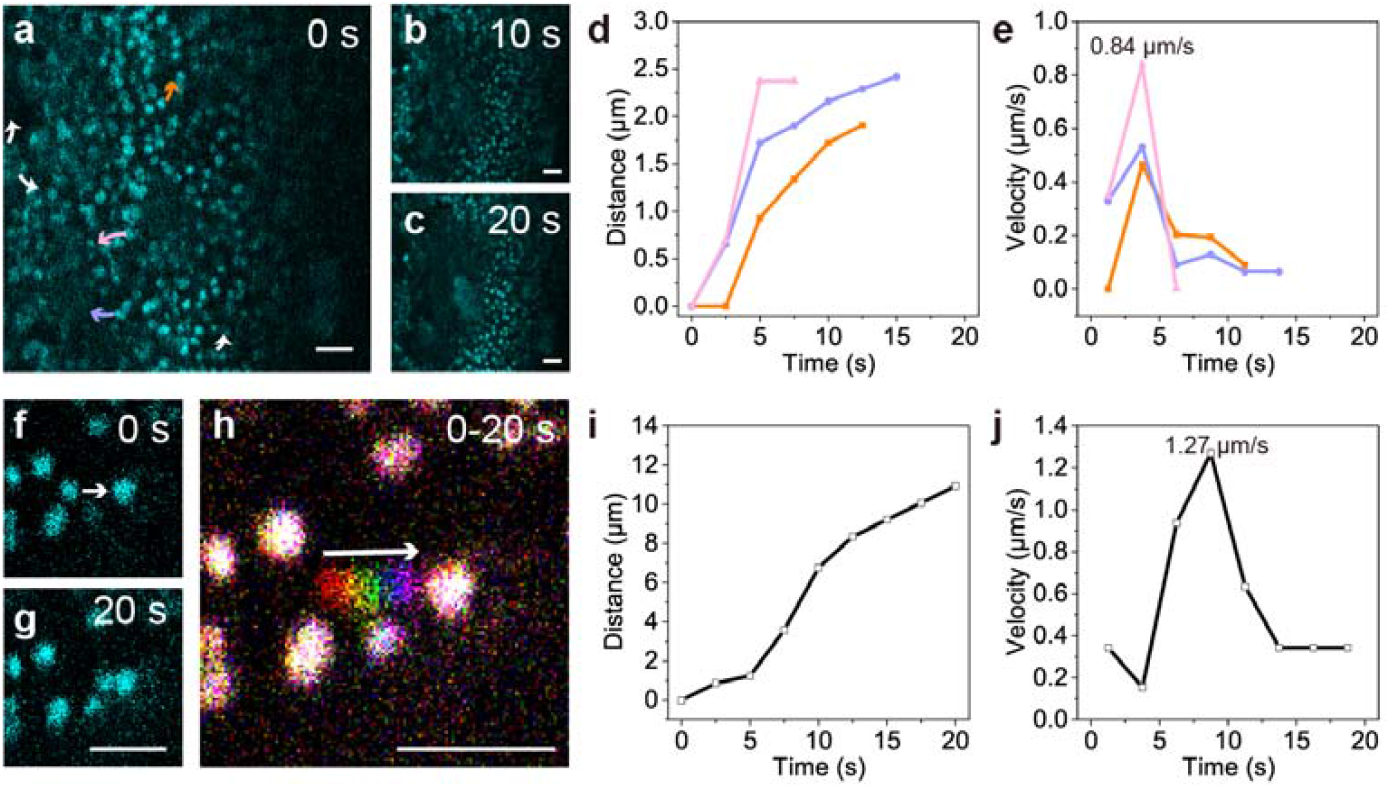
Quantitative measurement of rapid chloroplast avoidance movement in living submerged leaves of *Hydrocotyle sibthorpioides* by NIR-II fluorescence confocal microscope under 671 nm laser excitation with detection beyond 1000 nm. (a–c) Time-lapse NIR-II fluorescence confocal microscopic images at 0 s (a), 10 s (b), and 20 s (c) under continuous scanning of 671 nm laser showing chloroplast relocation away from the focal plane. Arrows indicate the lateral movements of representative chloroplasts (with the tails of arrows as the beginning points) undergoing avoidance. (d, e) Lateral displacement and velocity quantification for multiple chloroplasts in (a-c), revealing peak velocities top out at 0.84 μm/s. The curves represent the lateral movements of chloroplasts at the tail of arrows of the same color. (f, g) NIR-II fluorescence confocal microscopic images (0 s and 20 s) within intact leaf with lower activity than active leaves (detached and kept at room temperature for ∼30 minute), showing lateral displacement of individual chloroplast from the original positions. The images were enlarged to show the moving chloroplast. Arrow indicates the movement direction of the chloroplast. (h) Composite image pseudo-colored to illustrate the movement dynamics of chloroplast in (f, g) over 20 s. The arrow indicates the temporal sequence of different pseudo-colors. (i, j) Lateral displacement and velocity of an individual chloroplast in (h) showing a peak lateral velocity of 1.27 μm/s. Scale bars: 20 μm.

## Discussion

We established a noninvasive NIR-II fluorescence confocal microscope approach that enables real-time, three-dimensional visualization of chloroplast dynamics in intact leaves. By exploiting the fluorescence tail of chloroplasts in NIR-II spectral region and a single red laser beam as both the stimulation light and excitation light of imaging, we captured rapid avoidance responses in submerged leaves of *Hydrocotyle sibthorpioides* and *Hygrophila pinnatifida*. These findings overturn the prevailing notion that chloroplast relocation is inherently slow and provide an intuitive microscopic illustration of the motility and plasticity in chloroplast of amphibious plants. Our results provide the first microscopic evidence that environmental light regimes fundamentally shape chloroplast behavior, and establish NIR-II fluorescence confocal microscopy as a powerful tool for in situ plant photobiology.

Our study challenges the conventional understanding of chloroplast photorelocation. Previous reports, based largely on transmittance measurements or visible-light microscopy, characterized chloroplast migration as a gradual process occurring over minutes, with rates of only a few micrometers per minute. By contrast, in situ NIR-II fluorescence confocal microscopic imaging of leaves revealed that chloroplasts in submerged leaves can relocate on a second timescale, reaching micrometer-per-second velocities. This suggests that the actin-driven motility machinery can operate far more dynamically than previously assumed, enabling plants to rapidly adapt to sudden changes in illumination.

The discovery of distinct response patterns between aerial and submerged leaves underscores the strong environmental dependence of chloroplast motility. Submerged leaves, which develop under low red-light availability due to light absorption of water, exhibited heightened sensitivity and robust avoidance under red-light stimulation. Aerial leaves, in contrast, showed little avoidance and instead displayed fluorescence enhancement under identical red-light stimulation, possibly reflecting distinct photosynthetic optimization strategies. These contrasting responses provide direct visual evidence of adaptive plasticity, linking environmental light conditions to organelle-level photoprotection mechanisms.

These biological results demonstrate the utility of NIR-II fluorescence confocal microscopy for deep-tissue imaging in plant systems. Unlike two-photon fluorescence microscopy, which suffers from wavelength mismatch and phototoxicity, NIR-II fluorescence confocal microscopic imaging exploits the intrinsic fluorescence tail of chloroplasts to achieve subcellular resolution, extended imaging depth, and simultaneous stimulation-imaging capability with minimal perturbation—capabilities difficult to achieve with conventional visible fluorescence confocal or two-photon fluorescence microscopy. This opens avenues for real-time, in vivo studies of plant organelles that were previously inaccessible. This methodological advance provides a versatile platform for probing organelle dynamics deep within intact plant tissues. Future integration of NIR-II fluorescence confocal microscopy with molecular genetics will be essential to uncover the signaling and cytoskeletal mechanisms underlying rapid chloroplast motility. Extending this approach to other species and ecological settings will determine whether the fast responses of chloroplasts are widespread adaptations and how they contribute to plant photosynthesis in fluctuating light environments. More broadly, the ability to image rapid subcellular processes in large-scattering tissues of NIR-II fluorescence confocal microscopy opens new avenues for plant biology, bridging optical innovation with ecological and agricultural applications.

## Materials and Methods

### Plant Samples and Growth Conditions

*Hydrocotyle sibthorpioides* and *Hygrophila pinnatifida* were initially cultivated in moist plant substrate under controlled environmental conditions. The plants were exposed to LED lighting for 7 hours per day and maintained at a constant temperature of 25□°C.

To induce submergence acclimation, both *Hydrocotyle sibthorpioides* and *Hygrophila pinnatifida* were divided to retain part of aerial leaves, then transferred to containers filled with aquatic substrate and submerged. CO□ was supplied using a gas cylinder, and the same lighting and temperature conditions were maintained throughout the submergence period (7-hour daily LED illumination at 25□°C). Within approximately 2-4 days, aerial roots began to develop. Aerial leaves of *Hydrocotyle sibthorpioides* exhibited noticeable thinning, yellowing, and increased transparency, while those of *Hygrophila pinnatifida* developed dark spots and blackening along margins. After 7-10 days, most of the original aerial leaves had dissolved, and new leaves emerged, marking the completion of the submergence acclimation process. Once the plants reached a stable growth state underwater, leaves were collected for experiments.

### Visible Fluorescence Confocal Microscopic Imaging of Intact Leaf Samples

Visible fluorescence confocal microscopic imaging was performed using a commercial system (Zeiss LSM 900) with an objective (63x oil, NA 1.4). A single intact plant was selected, and individual leaves were flattened and fixed between a microscope slide and a coverslip, then placed on the microscope stage for imaging. The laser power in front of the objective was set to 1□mW for both 405□nm and 640□nm excitation. For time-lapse microscopic imaging in Video S2, a higher power setting of 6□mW in front of the objective was used for 405□nm excitation.

### NIR-II Fluorescence Confocal Microscopic System

A 671 nm CW laser beam was directed traversing into the microscope. A Dichroic Cage Cube (DM cube, MCM1S-A, LBTEK, China) housed an 1000 nm short pass dichroic mirror (DM10-1000SP, LBTEK, China). The excitation beam, after passing through this dichroic mirror, traversed the galvanometric mirrors, scan lens, tube lens, and was finally focused onto sample by a microscope objective (60x oil, NA 1.3, Olympus, Japan). The emitted NIR-II fluorescence was reflected by the short-pass dichroic mirror in the Dichroic Cage Cube. Then, the fluorescence signal passed through an interchangeable long-pass filter (EFLH1000, EFLH1100, EFLH1200, or EFLH1450, Thorlabs, USA), a Reflective collimator (RC02APC-P01, Thorlabs, USA), and coupled into a multimode fiber. The signal was then transmitted via the fiber to a photomultiplier tube (PMT, H12397A-75, Hamamatsu, Japan) for detection.

### NIR-II Fluorescence Confocal Microscopic Imaging of Intact Leaf Samples

For imaging of active leaves, a single intact plant was selected, and individual leaves were flattened and fixed between a microscope slide and a coverslip, then placed on the microscope stage for imaging. For metabolically inactive leaves, imaging was conducted 2 hours after leaf collected. To detect the lateral moving chloroplasts in figure 6, imaging was conducted 30 minutes after leaf collected. During standard imaging, the laser power in front of the objective was set to 1□mW. For high-power control experiments, the laser power in front of objective was increased to 4.5□mW. For low-power control experiments, the laser power in front of objective was reduced to 0.2□mW.

## Data Availability

All data generated or analyzed during the study are included in the articles and supplemental information published herein.

## Acknowledgments

This work was supported by the National Key R&D Program of China (2022YFB3206000), National Natural Science Foundation of China (61975172) and Hangzhou Chengxi Sci-tech Innovation Corridor Management Committee funded project.

## Author Contributions

J.Q. and W.G. conceived and designed the project. W.G. performed the preparation of materials and imaging experiments. M.H. ran the integrating spectra test. W.G. performed the imaging analysis and wrote the original manuscript. J.Q. and M.H. assisted with manuscript editing. J.Q. supervised the research.

## Conflict of Interest Statement

All authors disclosed no relevant relationships.

## Notes

### Competing Interest Statement

The authors have declared no competing interest.

